## Supplementary Materials for "Intraspecific and intraindividual trait variability decrease with tree species richness in a subtropical tree biodiversity experiment"

### SUPPORTING INFORMATION

**Fig. S1.** Regression coefficients for the effects of tree species richness on the intraspecific and intraindividual variability on seven leaf functional traits and two main axes of leaf trait variability.

**Fig. S2.** Results of a principal component analysis (PCA) of seven leaf functional traits belonging to eight different tree species.

**Fig. S3.** Location of eight sampled species in a functional trait space assessed by a principal component analysis (PCA) for seven leaf functional traits.

**Fig. S4.** Conceptual model representing the relationships between variables that could affect intraspecific overlap in leaf functional traits.

**Fig. S5.** Effect of tree species richness on intraspecific overlap.

**Fig. S6.** Results of non-simplified piecewise structural equation models (SEM) studying the mechanisms driving the intraspecific overlap in leaf functional traits.

**Fig. S7.** Conceptual framework for measuring community functional diversity based on individual leaf trait values (following the approach of Carmona et al. 2016 (37)).

**Fig. S8.** Conceptual framework for the null model approach based on the randomization of different sources of variation.

**Fig. S9.** Results of linear mixed-effects models to test the joint effect of tree species richness and the type of null models on standardized effect sizes (SES) of two univariate functional indices (functional richness (FRic) and functional divergence (FDiv)) calculated from the two main axes of leaf variation (PC1 and PC2) and for four different sources of trait variation.

**Fig. S10.** Differences in stomatal density across different positions in the tree crown.

**Fig. S11.** Bar plots for the variance partitioning of leaf variation.

**Fig. S12.** Location and slope of the sampled trees within the experimental site.

**Fig. S13.** Leaf reflectance spectra for the eight study species.

**Fig. S14.** Scatter plot of predicted and measured trait values in the test and the train samples.

**Fig. S15.** Evolution of the error during the training of convolutional neural networks for trait prediction.

**Fig. S16.** Bar plot and heatmap of the distribution of missing trait data in the leaf-level dataset.

**Fig. S17.** Excluded values from predicted leaf-level data for seven leaf functional traits.

**Fig. S18.** Analytical framework used to assess the metrics of intraindividual variability, intraspecific variability and intraspecific overlap.

**Fig. S19.** Evolution of mean and variance of simulated values of FRic and FDiv from different null models with increasing number of randomizations.

**Table. S1.** Summary of a principal component analysis for seven leaf functional traits, including loadings, standard deviation, proportion of the variance explained by each component and the adjusted eigenvalue obtained in a Horn's parallel analysis.

**Table. S2.** Results for linear mixed-effects models studying the effects of tree species richness on multivariate functional indices used to estimate intraspecific variability, intraindividual variability and intraspecific overlap.

**Table. S3.** Species included in the study.

**Table. S4.** Results for linear mixed-effects models studying the effects of tree species richness and type of null model on standardized effect sizes of two functional indices.

**Table. S5.** Layers and hyperparameters used for building a convolutional neural network for every trait, and coefficient of determination ($R^2$) and root mean squared error (RMSE) for the test and the train samples.

**Table. S6.** Distribution of missing trait data in the leaf-level dataset across species and traits.

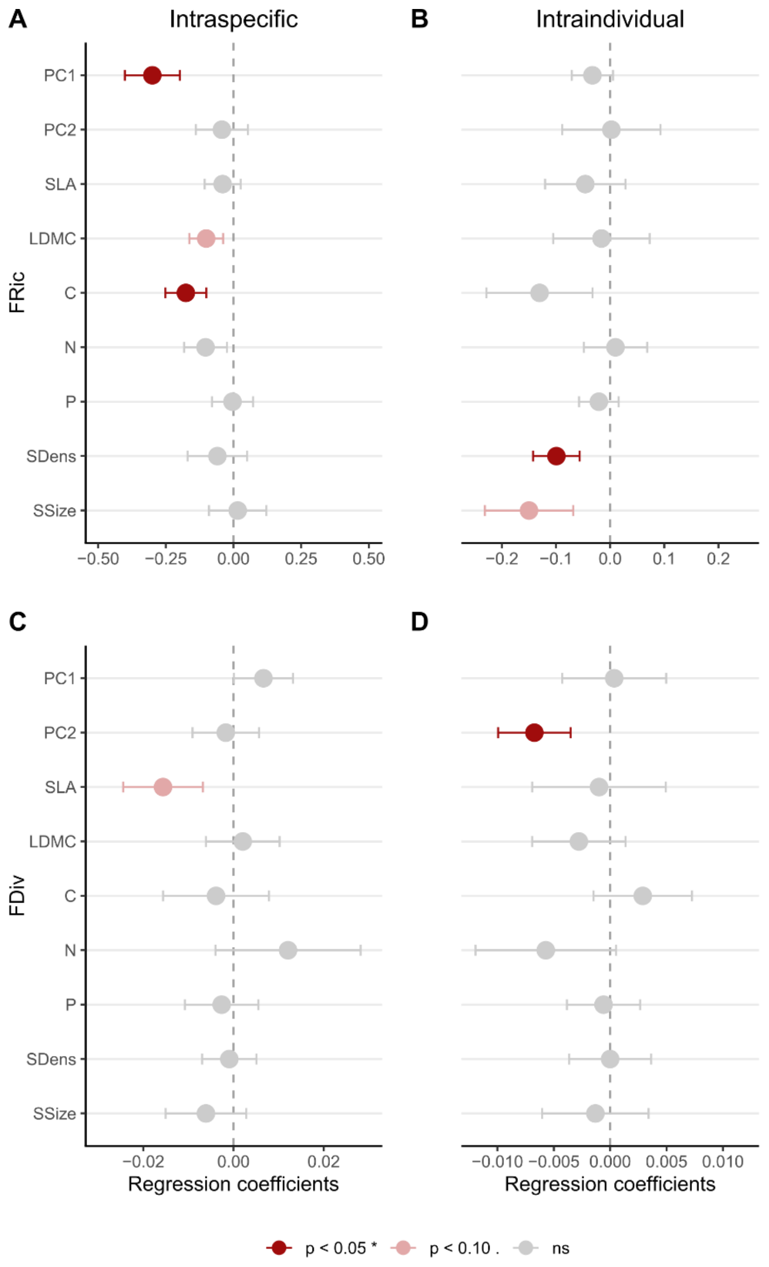


**Fig. S1. Regression coefficients for the effects of tree species richness on the intraspecific and intraindividual variability on seven leaf functional traits and two main axes of leaf trait variability.** Regression coefficients for the effects of tree species richness on the intraspecific and intraindividual variability on seven leaf functional traits and two main axes of leaf trait variability. The effects of tree species richness on intraspecific and intraindividual variability were studied for seven functional traits (specific leaf area (SLA), leaf dry matter content (LDMC), leaf carbon content (C), leaf nitrogen content (N), leaf phosphorus content (P), stomatal density (SDens) and stomatal size (SSize)) and for two main axes of leaf trait variability (PC1 and PC2). Colors represent the significance as determined by a likelihood ratio test (red $p < 0.05$, pink $< 0.01$, grey $p > 0.05$).

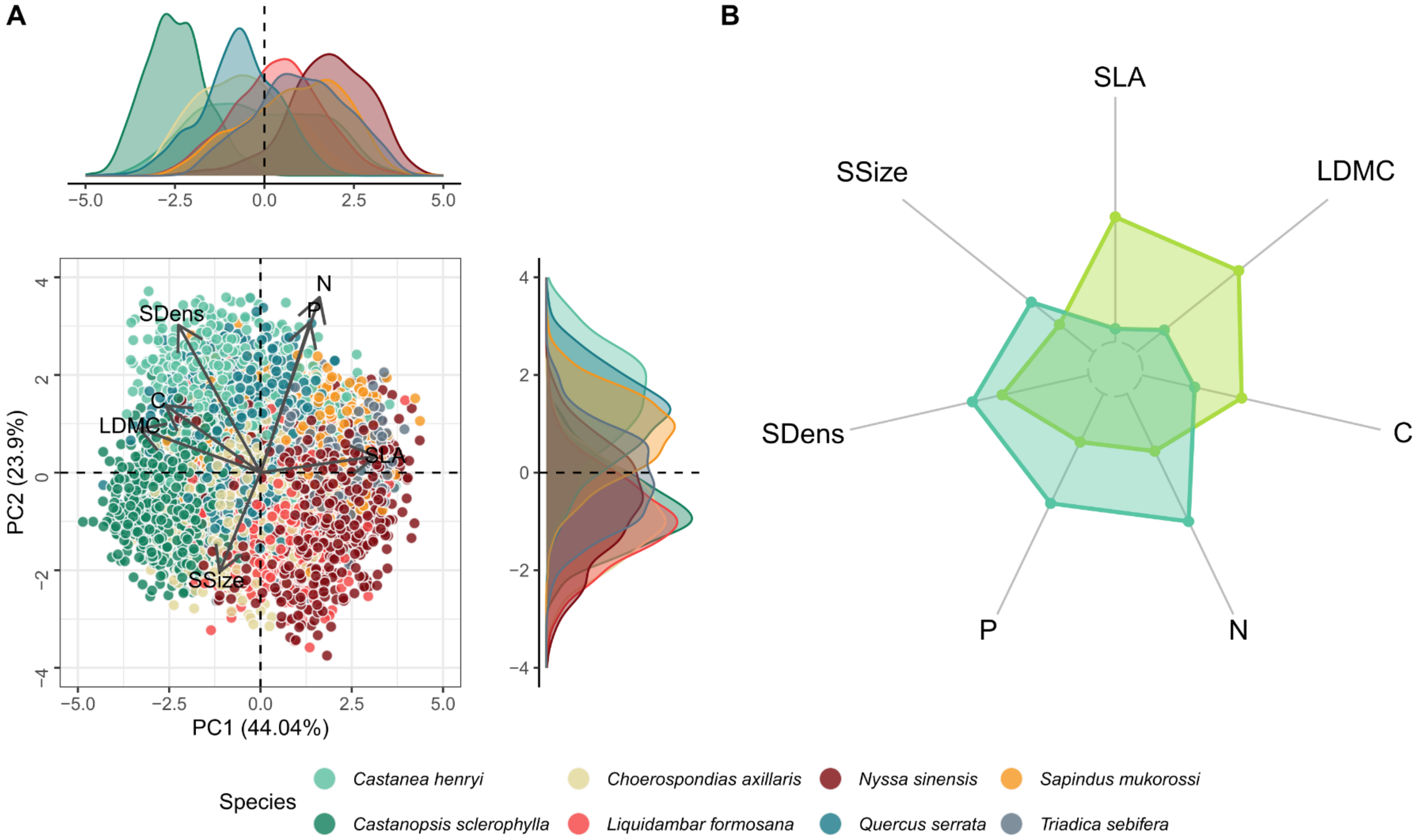


**Fig. S2. Results of a principal component analysis (PCA) of seven leaf functional traits belonging to eight different tree species.** (A) Main axes of a principal component analysis, including the location for every leaf and arrows representing the eigenvalues of every trait in the PCA axes, and (B) radar plot representing the eigenvalues of the traits in the two main axes. The first axis represents the variation in growth strategy, with lower values associated with a conservative strategy while higher values correspond to an acquisitive strategy. The second axis mainly associates with stomata density (SDens) and leaf P and N. This suggests that higher evapotranspiration rates linked to higher SDens might be associated with a higher content of P and N, which are key nutrients for photosynthetic activity (1).

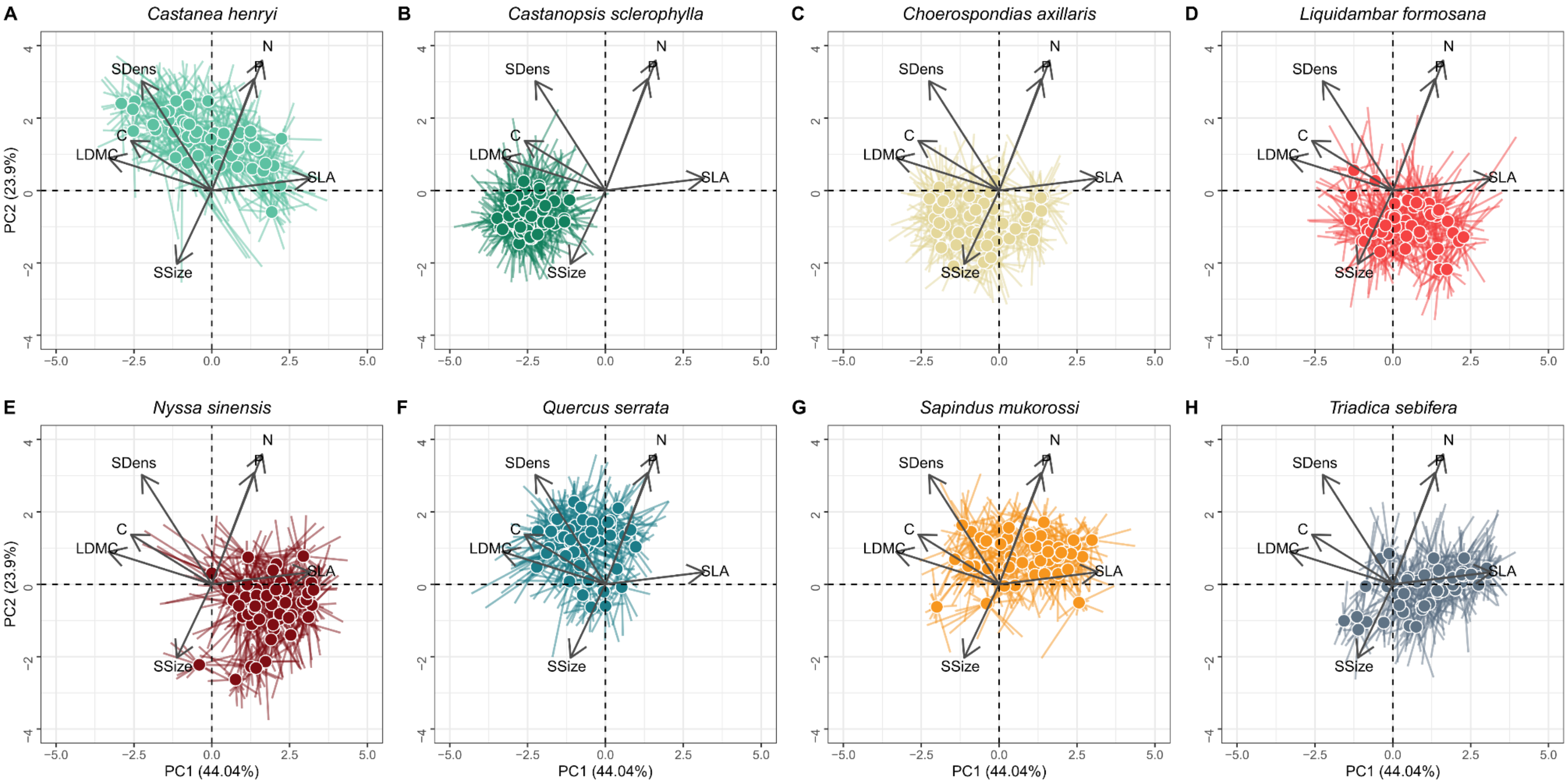


**Fig. S3. Location of eight sampled species in a functional trait space assessed by a principal component analysis (PCA) for seven leaf functional traits.** Each panel corresponds to a different species, dots represent the mean trait values of every tree (centroid) and lines connect the centroid to the individual values of the leaves belonging to the tree. The figure suggests that there are differences between species in terms of their leaf strategy (e.g. Castanopsis sclerophylla has a more conservative growth strategy compared to Triadica sebifera) and in terms of the extent of their intraspecific variation (e.g. Nyssa sinensis and Castanea henryi show high intraspecific variability compared to other species).

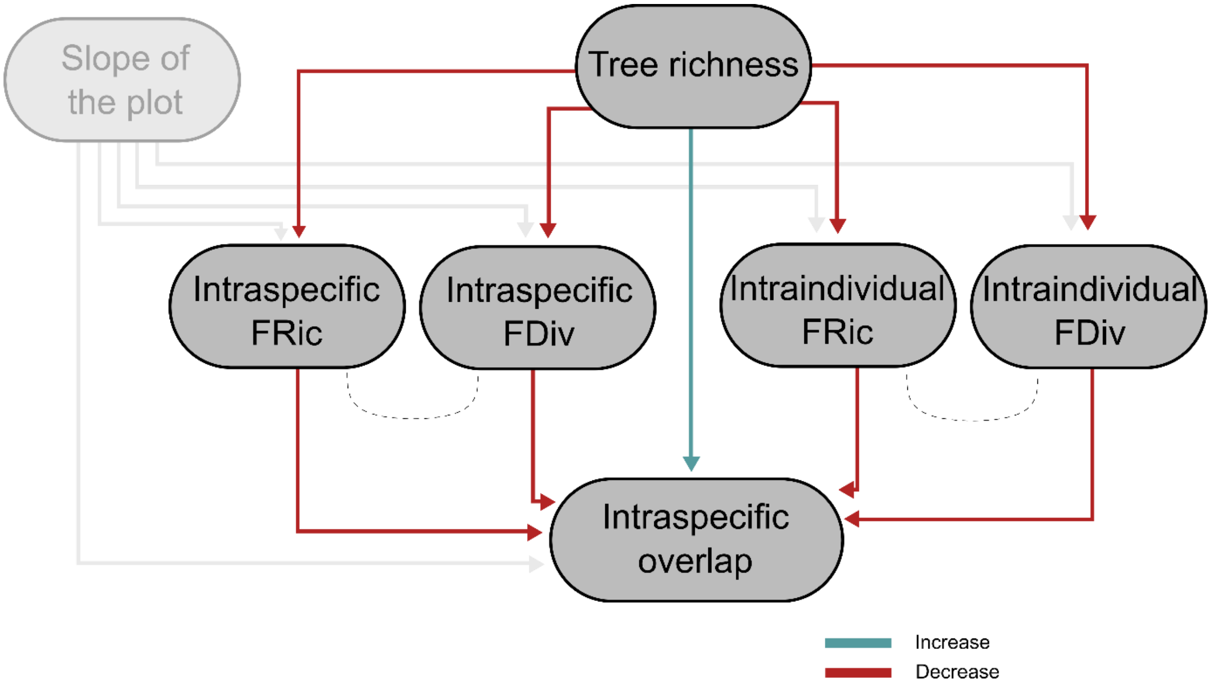


**Fig. S4. Conceptual model representing the relationships between variables that could affect intraspecific overlap in leaf functional traits.** Tree species richness is expected to affect negatively the intraspecific and intraindividual trait variability (for both indices), while these are expected to have a negative effect on the intraspecific overlap. This was expected as intraindividual and intraspecific trait variability were hypothesized to act as mechanisms to guarantee complementarity in intraspecific interactions. The slope of the plot was included as a covariate in the analyses in order to control for it, but hypotheses were not formulated for its effect on the other variables. Red lines represent negative relationships, blue lines represent positive relationships and dashed lines indicate correlated error terms.

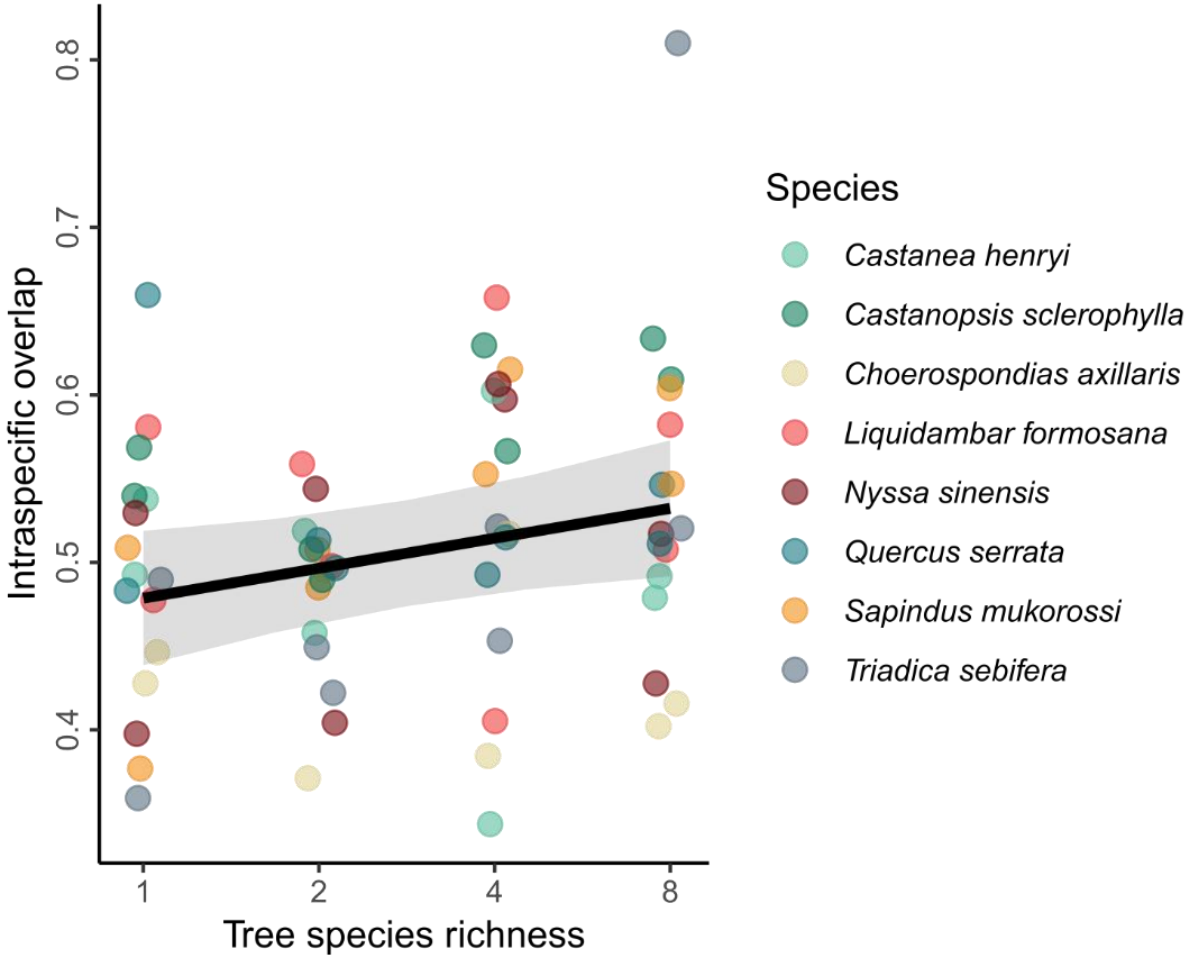


**Fig. S5. Effect of tree species richness on intraspecific overlap.** The line corresponds to the results of a linear mixed-effects model that shows a significant increase of intraspecific overlap with increasing tree species richness (P = 0.03, N = 63). Grey bands represent a 95% confidence interval. Colors correspond to the different tree species included in the study, whose identity was included as a random effect in our models. The slope of the terrain was included as a covariate in the model.

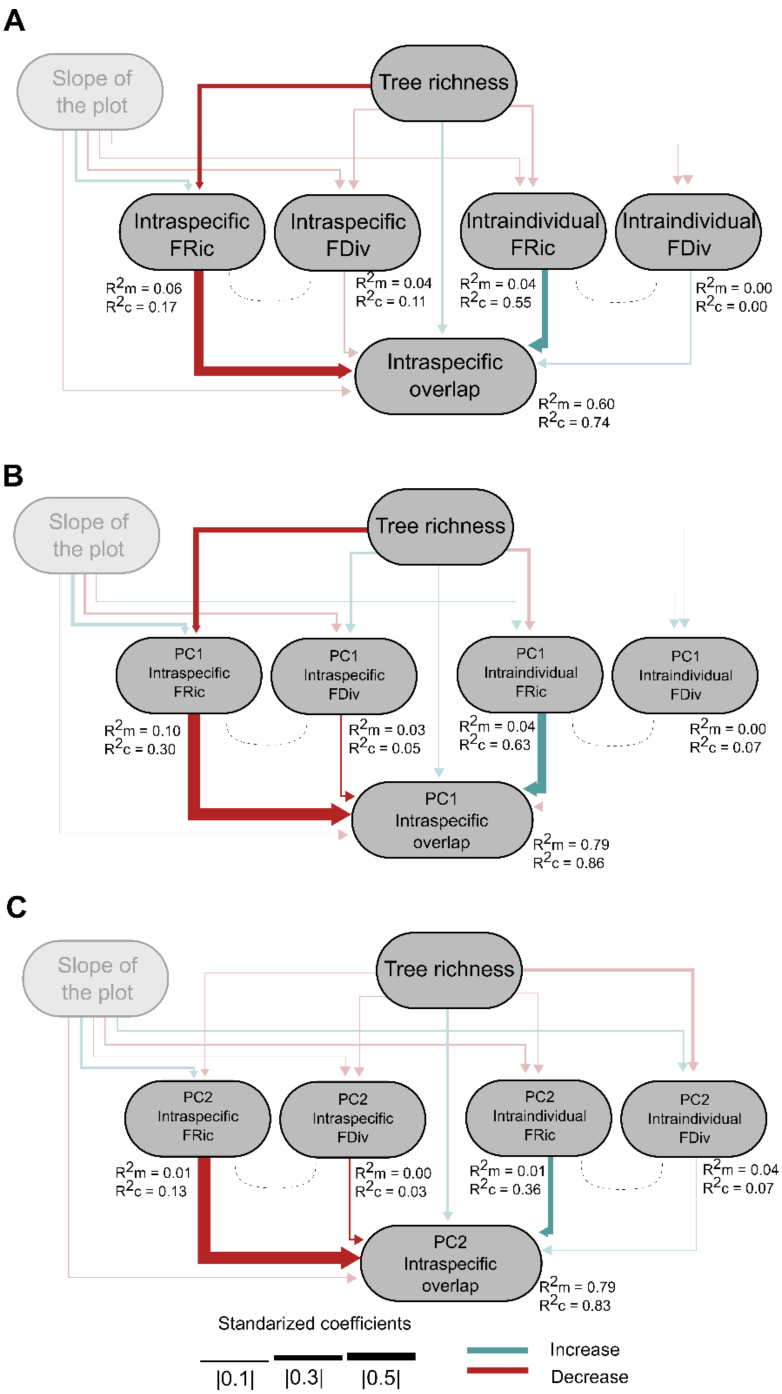


**Fig. S6. Results of non-simplified piecewise structural equation models (SEM) studying the mechanisms driving the intraspecific overlap in leaf functional traits.** Results are shown for (A) a complete SEM based on the conceptual model defined in Fig. S4, and for SEMs based on the variability on the two main axes of trait variation: (B) PC1 and (C) PC2. The width and color of the arrows indicate the strength and direction of the effects. Significant results are represented by solid lines while non-significant relationships are represented by semi-transparent lines. The marginal and conditional R2 are indicated for every model of the piecewise SEM.

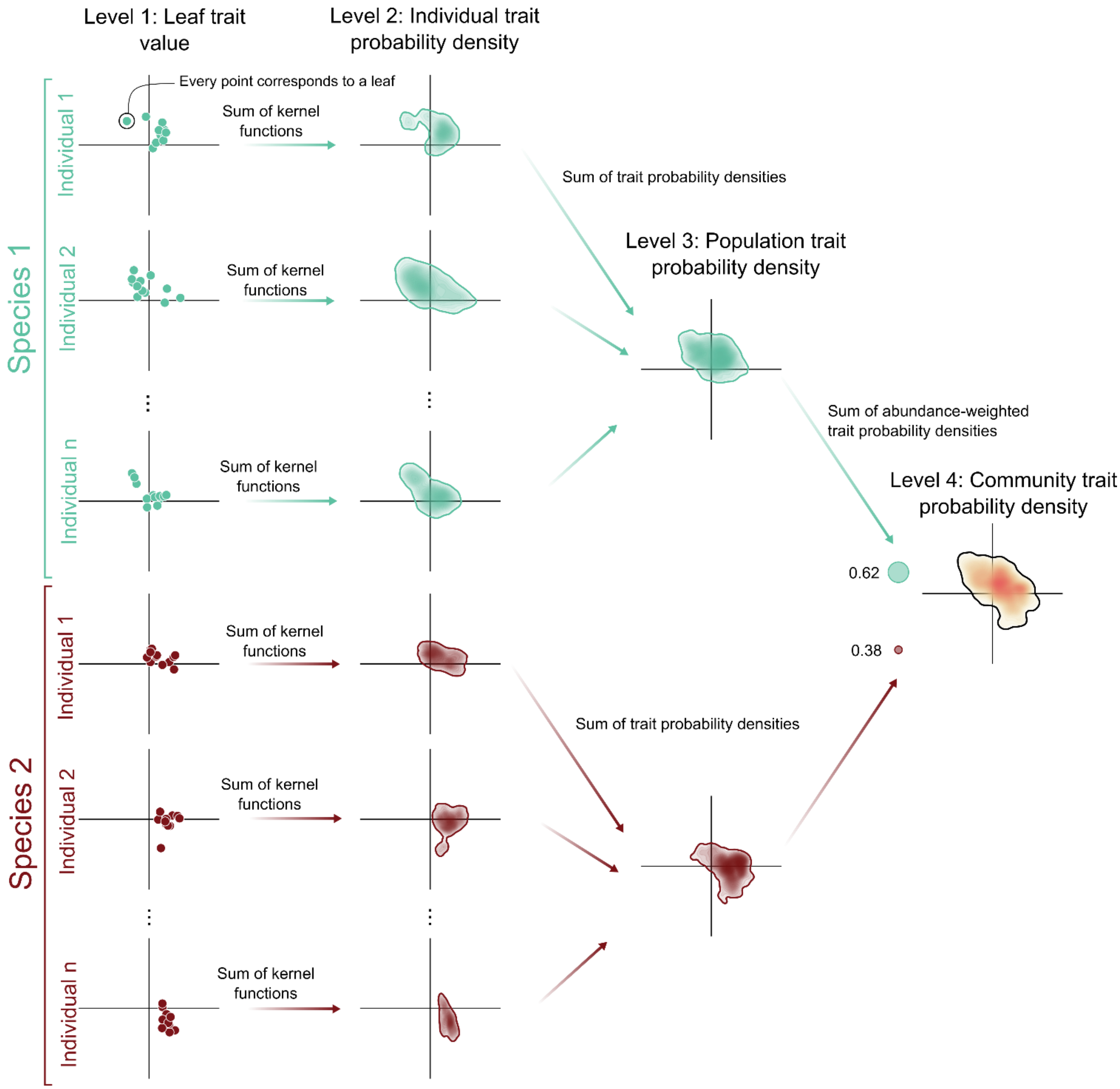


**Fig. S7. Conceptual framework for measuring community functional diversity based on individual leaf trait values (following the approach of Carmona et al. 2016 (37)).** As an example, we used a community with two species (species 1 in green and species 2 in red). First (Level 1), we applied kernel density functions to the leaf trait values (represented by the leaf's position on axis 1 and axis 2 of a PCA; Fig. 3) of every tree. Next, we summarized the kernel density functions to get one trait probability density for every tree (Level 2). After this, we calculated the sum of the trait probability densities of all trees belonging to the same population to get the population trait probability density (Level 3). Finally, we aggregated the population trait probability densities (Level 4). In this step, each trait probability density was rescaled according to the relative abundance of the species in the community.

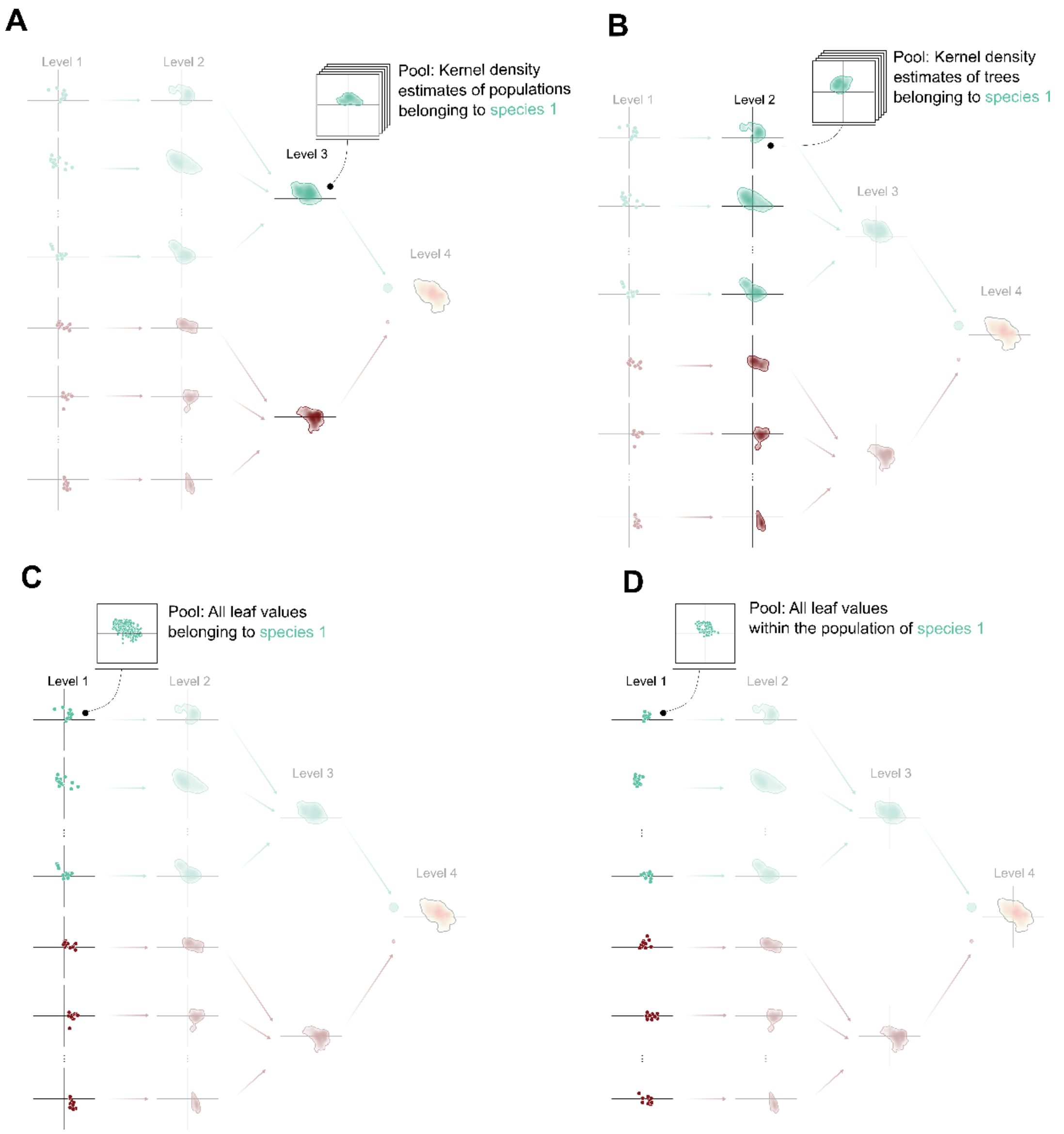


**Fig. S8. Conceptual framework for the null model approach based on the randomization of different sources of variation.** Null models differ in the process for generating simulated communities by randomizing different steps of the framework for measuring functional diversity as shown in Fig. S7. The result in every case is an assemblage with the same species composition and abundances as the observed one, but different levels of the variability occurring within the species were randomized. (A) The random population null model is generated by randomizing the population trait probability densities generated in step 3 by using as a pool all the different population trait probability densities calculated for that species. (B) The random tree null model is generated by randomizing the tree trait probability densities generated in step 2 by using as a pool all the different tree trait probability densities calculated for that species in any community. (C) The random leaf null model is generated by randomizing the leaves that are used to estimate the trait probability densities of trees by using as a pool all the different leaves for that species across the whole experiment. Finally, (D) The population-restricted random leaf null model is generated by randomizing the leaves that are used to estimate the trait probability densities of trees by using as a pool all leaves for that species within the population.

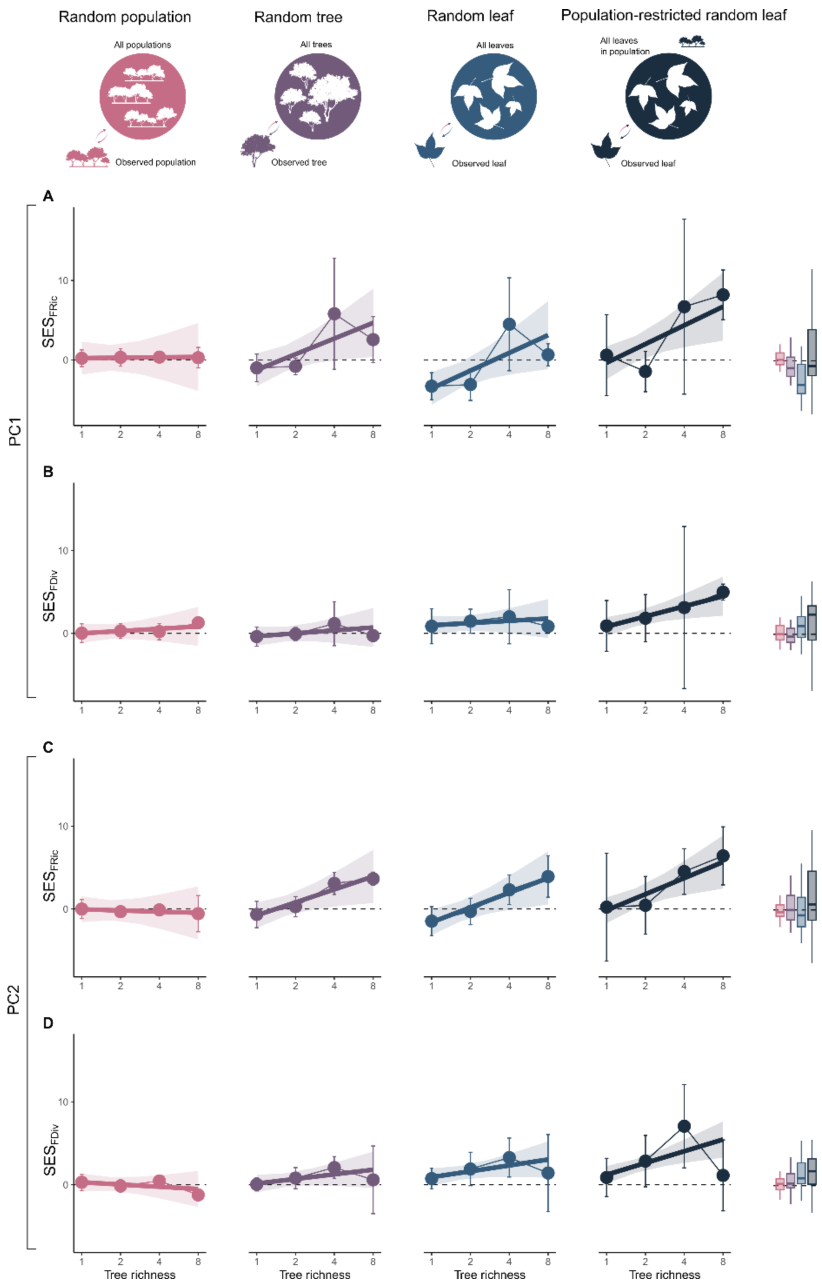


**Fig. S9. Results of linear mixed-effects models to test the joint effect of tree species richness and the type of null models on standardized effect sizes (SES) of two univariate functional indices (functional richness (FRic) and functional divergence (FDiv)) calculated from the two main axes of leaf variation (PC1 and PC2) and for four different sources of trait variation.** Linear mixed-effects models show a significant effect of the interaction of tree species richness and the type of model on SES(FRic) in the case of both axes ($P = 0.002$ for PC1 and $P < 0.001$ for PC1) and a significant effect of this interaction in the cases of SES(FDiv) in PC2 ($P < 0.001$), but this effect was not significant in the case of the SES(FDiv) of PC1 ($P = 0.17$). However, in the case of SES(FDiv) of PC1, the effects of tree species richness and the type of model were still significant ($P < 0.001$ in both cases).

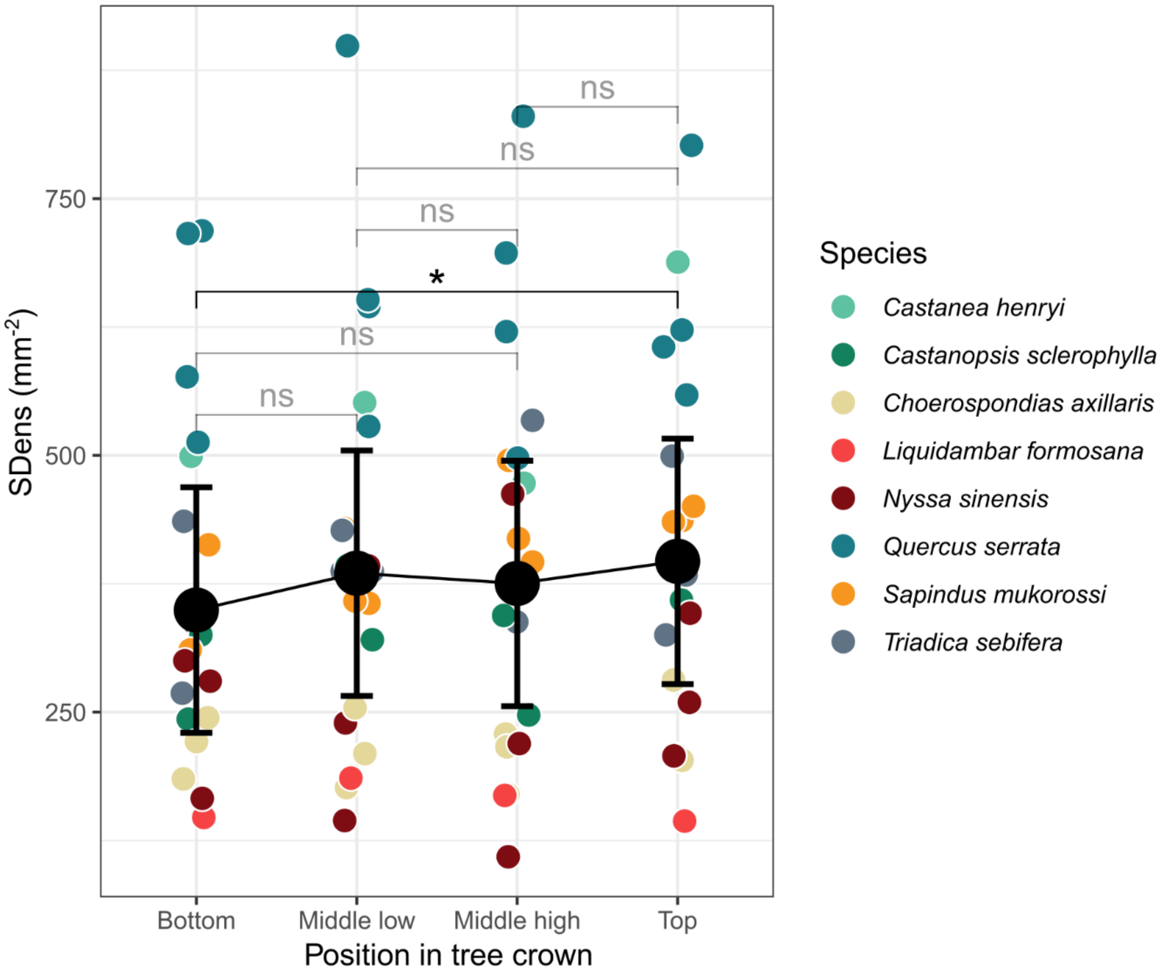


**Fig. S10. Differences in stomatal density across different positions in the tree crown.** Leaves of the calibration stomata set (see `Field sampling` section for description of the calibration stomata) were collected at different heights within the tree crown (bottom, middle low, middle high and top). A linear mixed-effects model to study differences in stomatal density (SDens) across different crown positions revealed significant differences between the top and the bottom ($P = 0.03$ as revealed by a Tukey post-hoc text). Black points and error bars represent estimate and confidence intervals, respectively. Every tree species is represented in a different color.

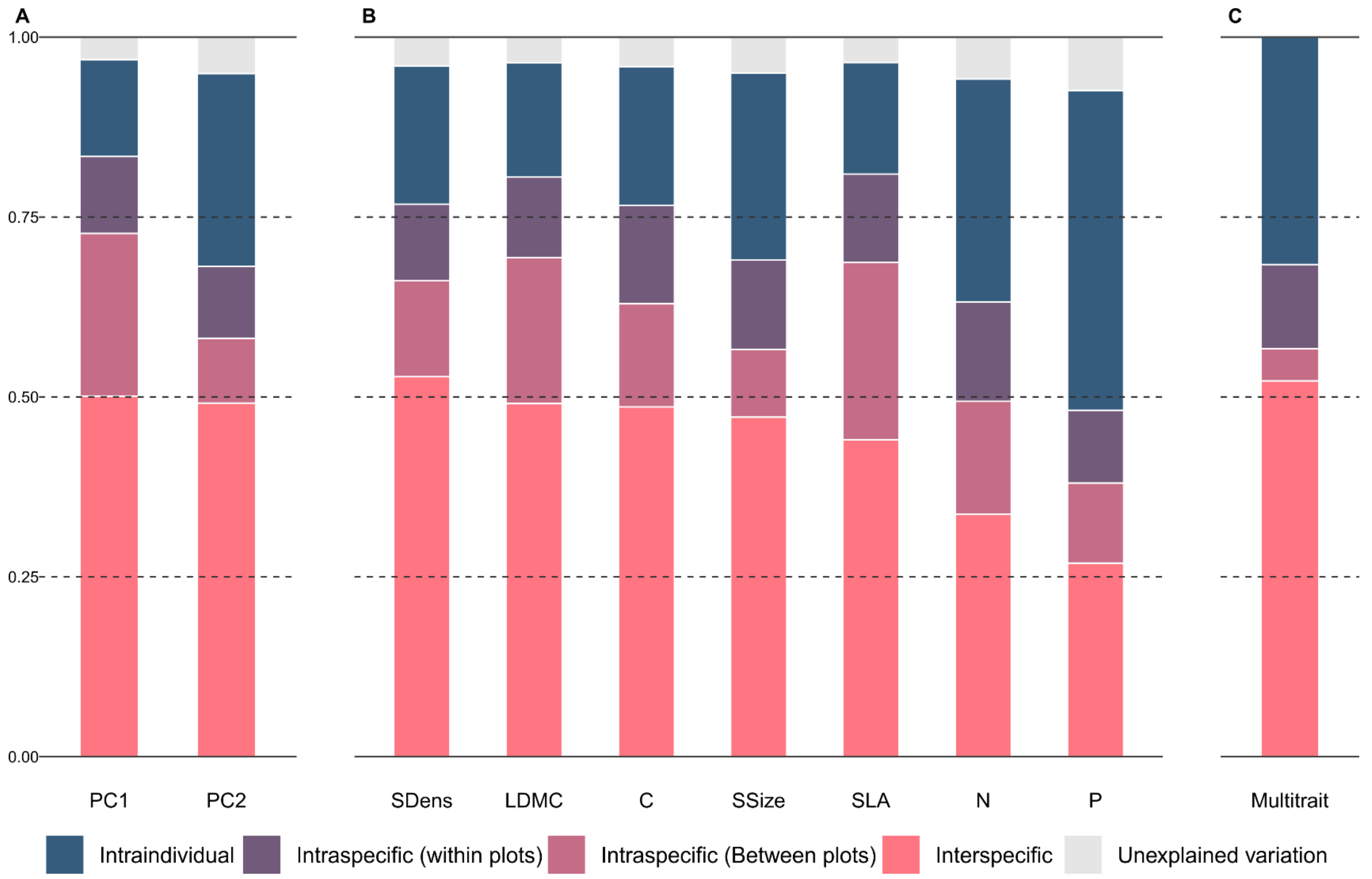


**Fig. S11. Bar plots for the variance partitioning of leaf variation.** Variance partitioning was studied for (**A**) the main axes of leaf variation found in a principal component analyses (PC1 and PC2; Fig. 2A), (**B**) independently for seven functional traits related to plant resource and water use (specific leaf area, SLA; leaf dry matter content, LDMC; leaf carbon content, C; leaf nitrogen content, N; leaf phosphorus content, P; stomatal density, SDens; stomatal size, SSize) and (**C**) jointly for the seven leaf traits mentioned. Variance partitioning in (**A**) and (**B**) was assessed by using an intercept only linear mixed-effects model with only random effects (leaf nested in tree, in turn nested in population, in turn nested in species identity), while we used a permutational multivariate analysis of variance (PERMANOVA) with the nested structure described as a predictor for the variance partitioning in (**C**).

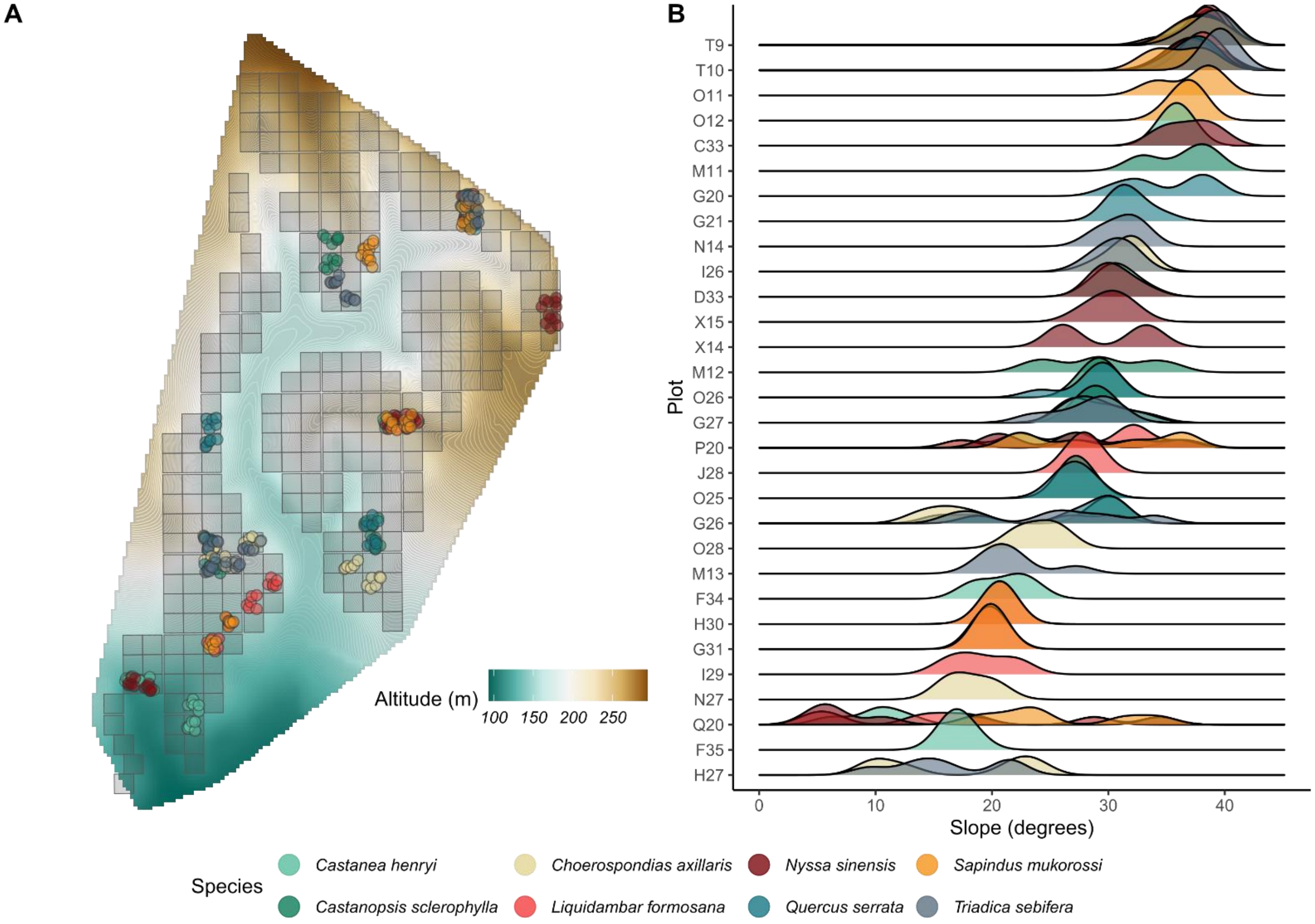


**Fig. S12. Location and slope of the sampled trees within the experimental site.** (**A**) Trees were sampled across 30 plots distributed in different parts of the experiment. (**B**) Density plots of the slope of every species in every plot (based on interpolated values of the slope of the terrain obtained from a 5 m resolution digital elevation model available at https://data.botanik.uni-halle.de/bef-china/datasets/53).

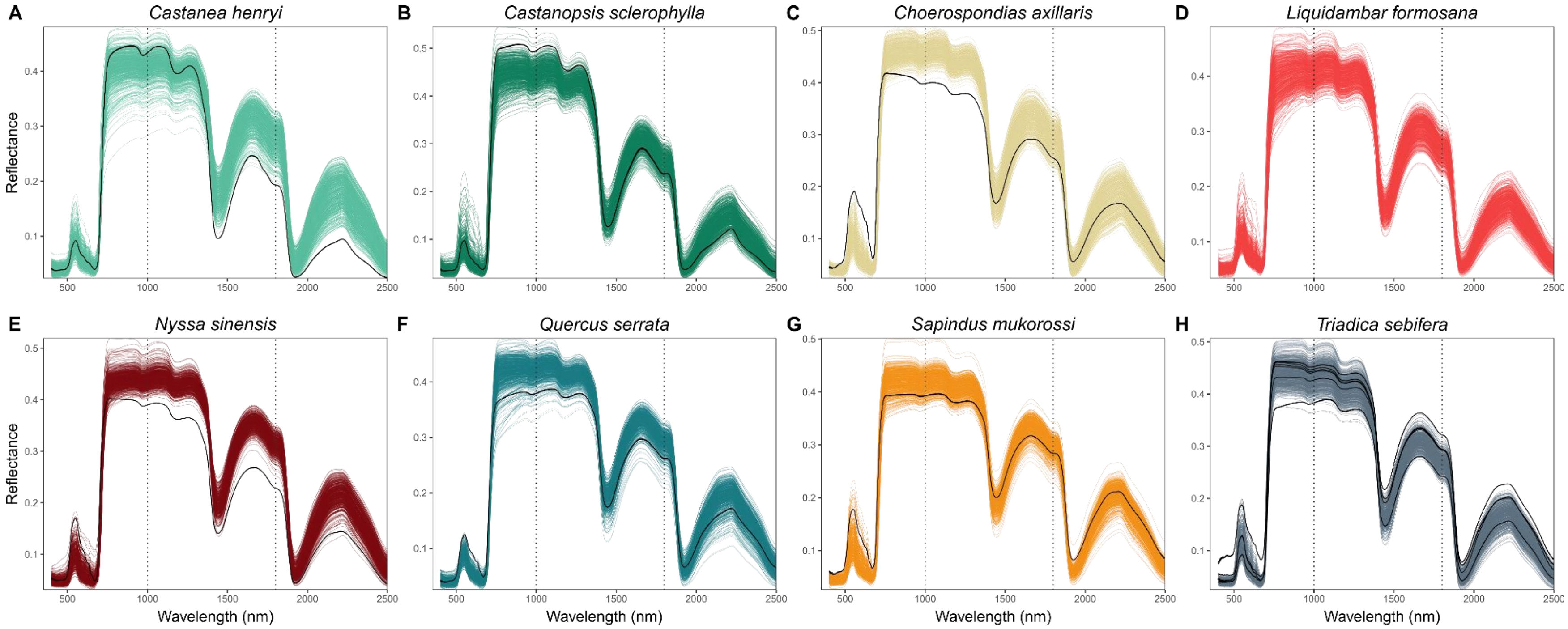


**Fig. S13. Leaf reflectance spectra for the eight study species.** Spectra of all leaves collected by species (represented in different panels and different colors). Lines in black represent those spectra, which were excluded for subsequent analyses as they had a local outlier factor higher than two. Dotted vertical lines represent the limits between the sensors of the spectroradiometer use.

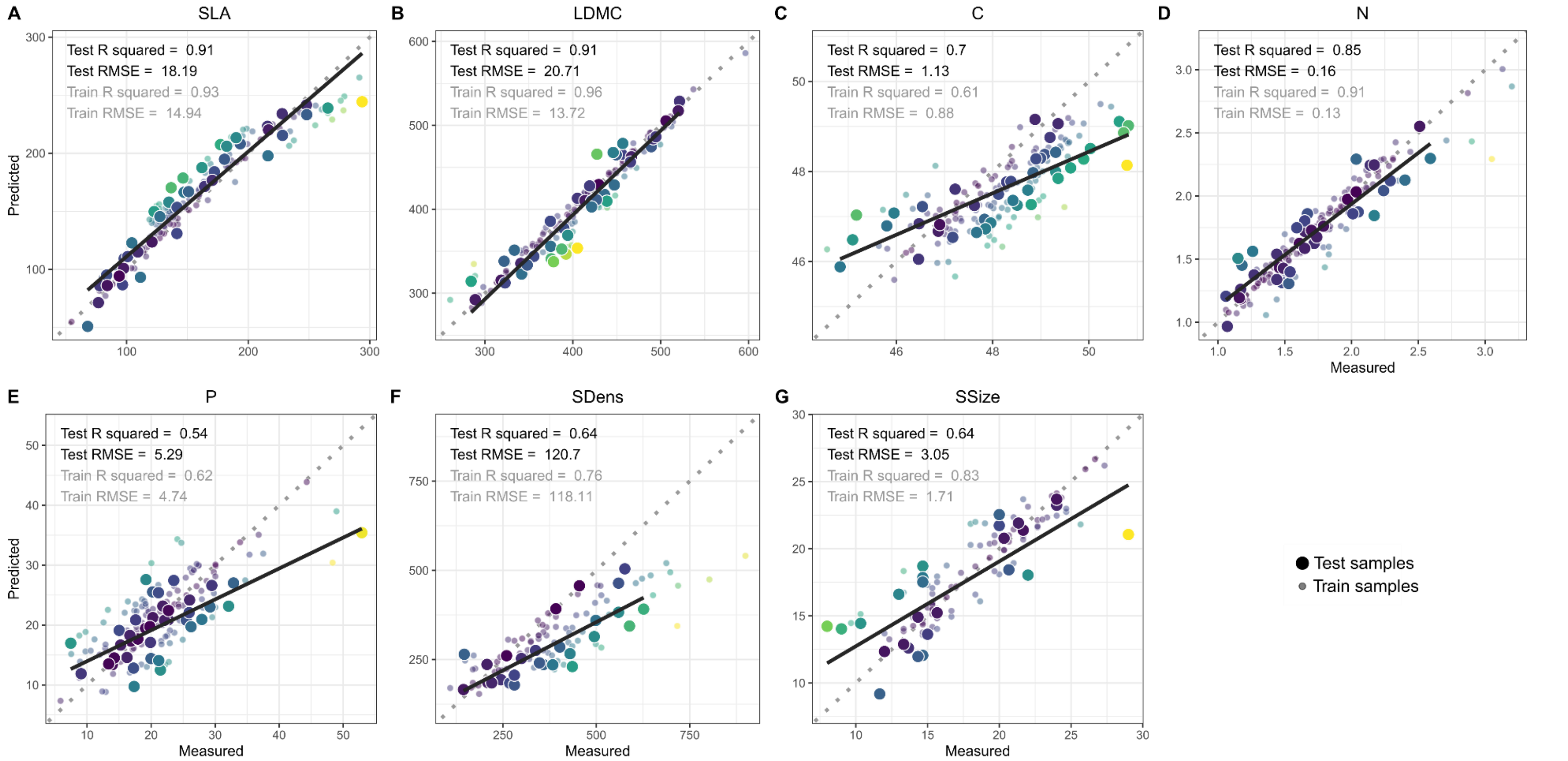


**Fig. S14. Scatter plot of predicted and measured trait values in the test and the train samples.** Correlation lines correspond only to the correlation between the predicted and measured values in the test samples. The dashed grey lines indicate the optimal fit in every case. The color of the points corresponds to the distance of the value from the optimal fit (dark blue colors indicate short distance while yellowish colors correspond to higher distances).

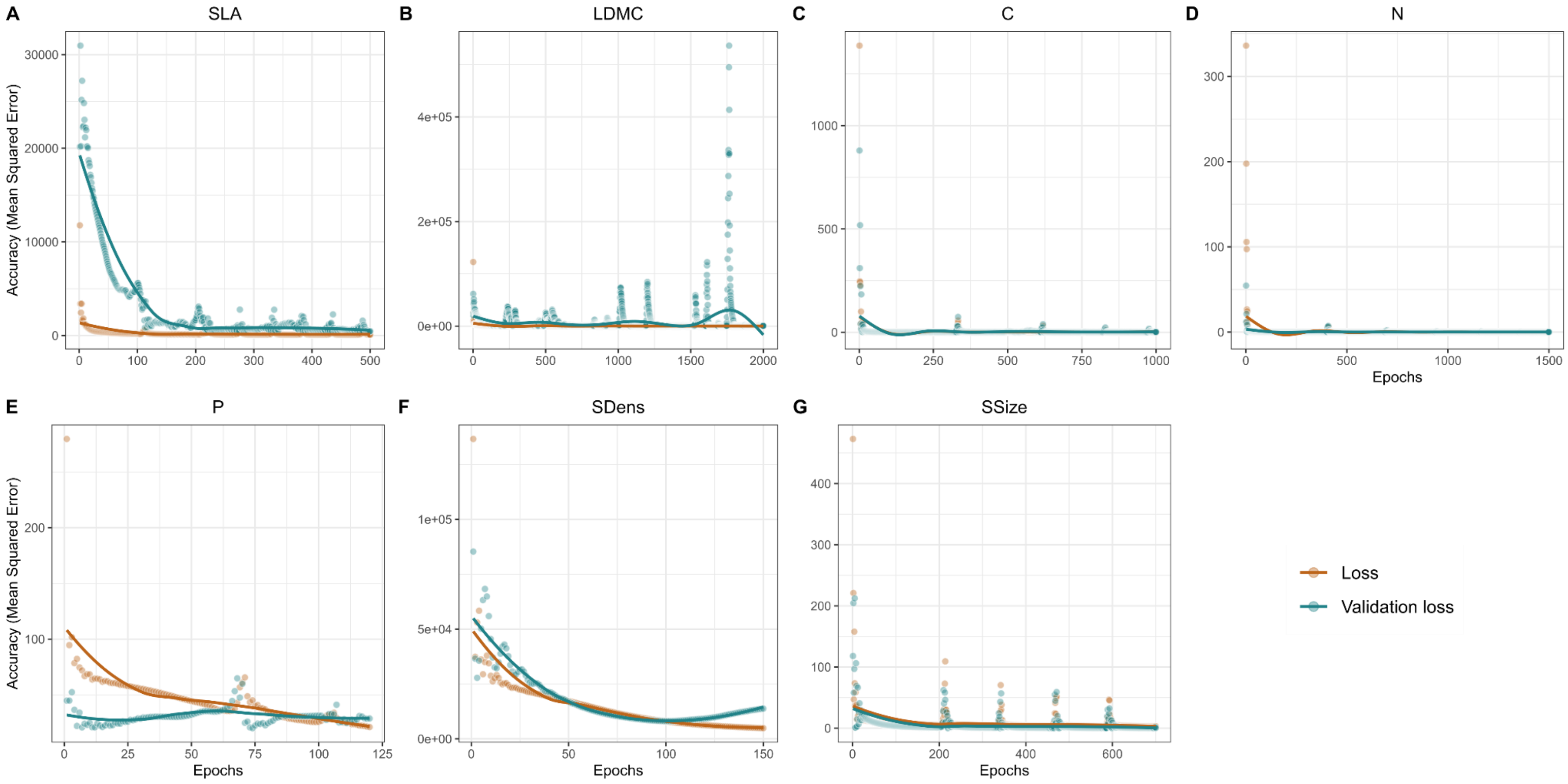


**Fig. S15. Evolution of the error during the training of convolutional neural networks for trait prediction.** Changes in the mean squared error in the loss function for all samples in the training set (in brown) and for a subset of samples used for validation during the training (in blue) were registered with an increasing number of epochs during the training process of the convolutional neural networks.

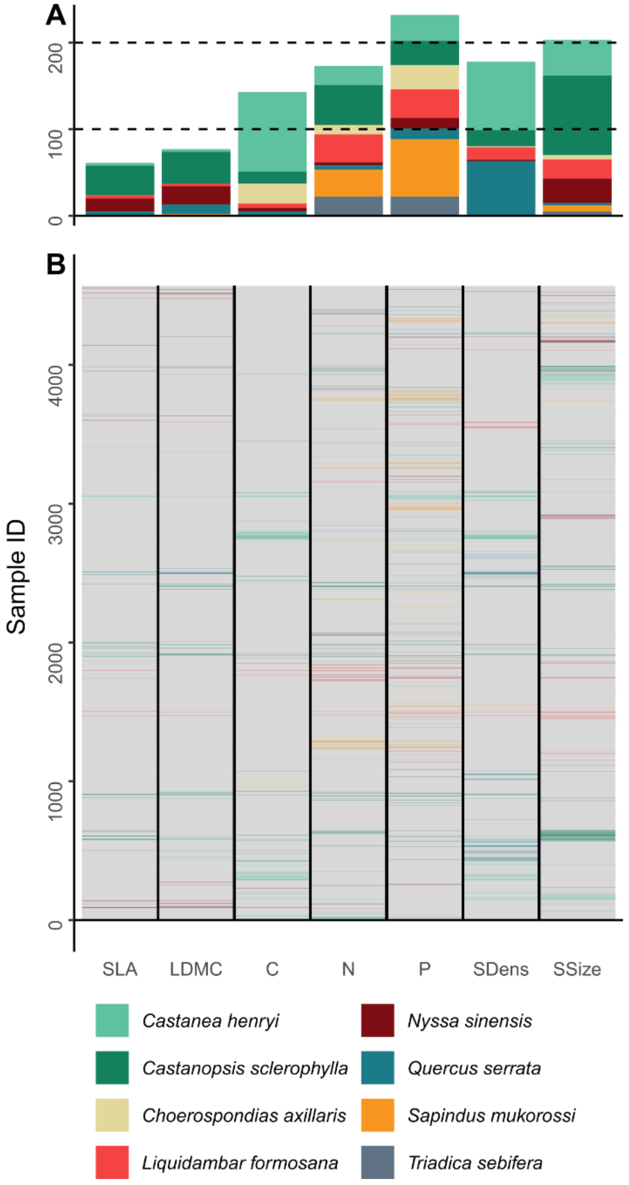


**Fig. S16. Bar plot and heatmap of the distribution of missing trait data in the leaf-level dataset.** (**A**) Represents a bar plot for the number of missing values for every trait (colored by species). In (**B**), vertical colored lines represent missing values and their distribution across the dataset (as ordered in the original dataset).

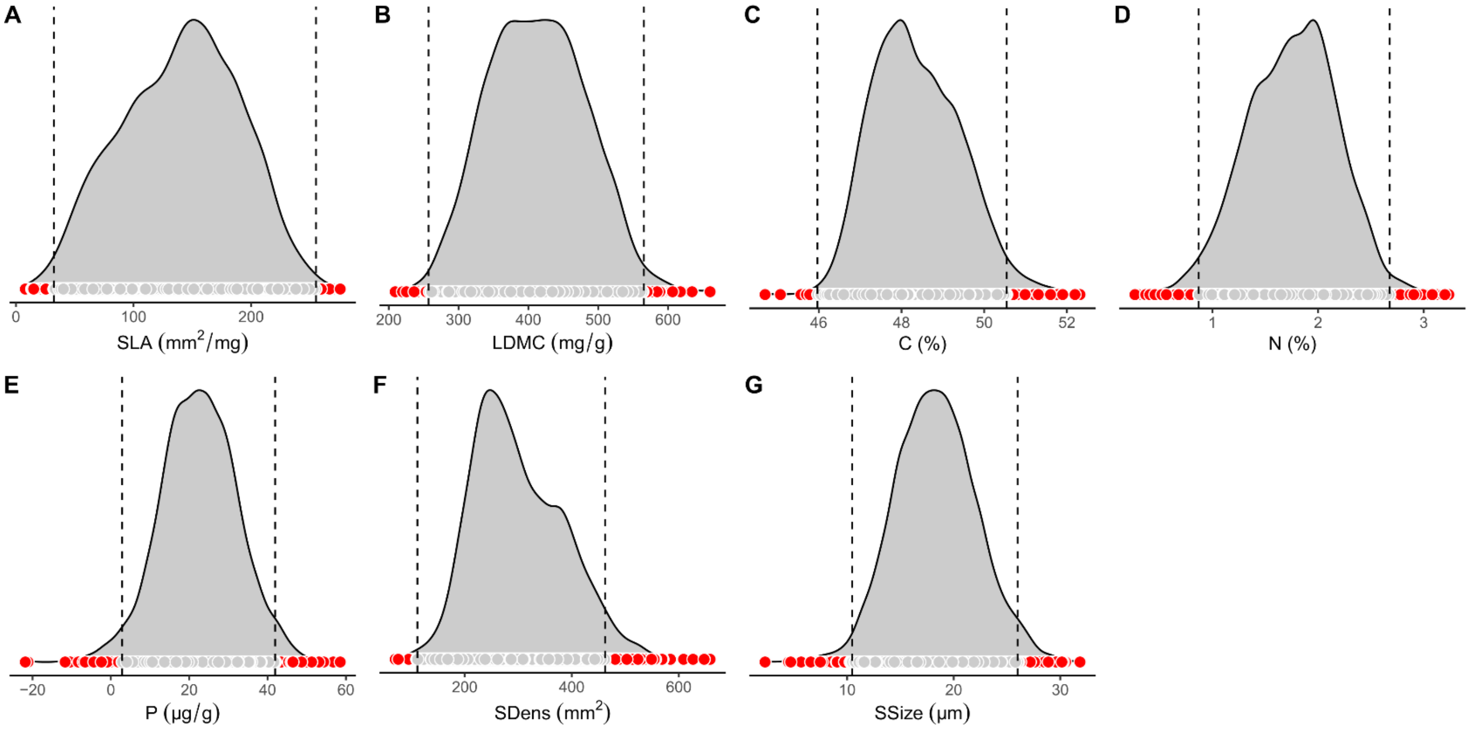


**Fig. S17. Excluded values from predicted leaf-level data for seven leaf functional traits.** Data excluded in trait predictions (showed in red) laid outside the interval formed by the median, plus or minus 3 median absolute deviations, as represented by the dashed vertical lines.

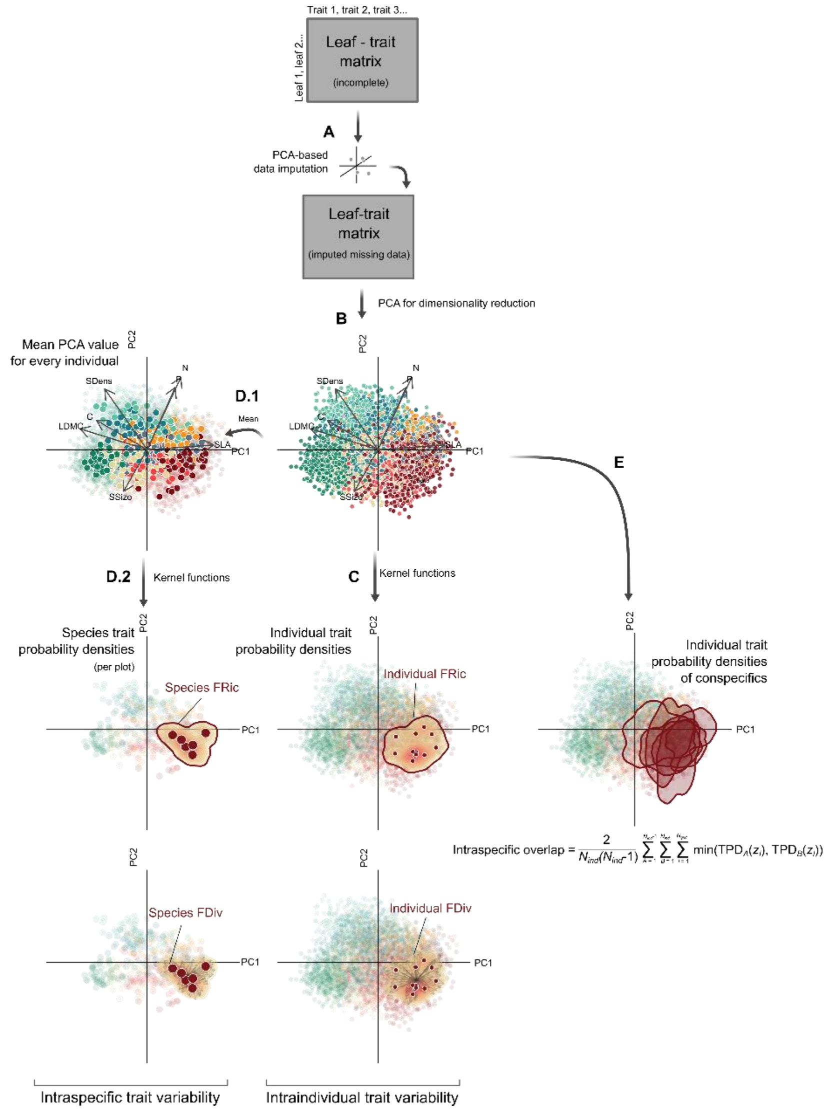


**Fig. S18. Analytical framework used to assess the metrics of intraindividual variability, intraspecific variability and intraspecific overlap.** All metrics were assessed by using (**A**) the leaf-level trait matrix. Due to missing values in the matrix, a principal component analyses (PCA)-based imputation approach was used to predict the missing data from the existing ones. With the completed dataset via imputation, (**B**) we performed a PCA to reduce the dimensionality of our data and used the first two principal components which together explained almost 70% of the variation. While, (**C**) trait probability densities were estimated for individual trees from this data, (**D.1**) mean values were obtained for individual trees in order to assess (**D.2**) trait probability densities for the intraspecific trait variability. From all these trait probability densities (the ones at the individual level and the population level) functional richness and functional divergence were used to estimate intraindividual and intraspecific trait variability. Last, (**E**) the trait probability densities estimated at the individual level for conspecifics (individuals from the same species occurring in the same plot) were used to estimate the mean intraspecific overlap of a species in a plot.

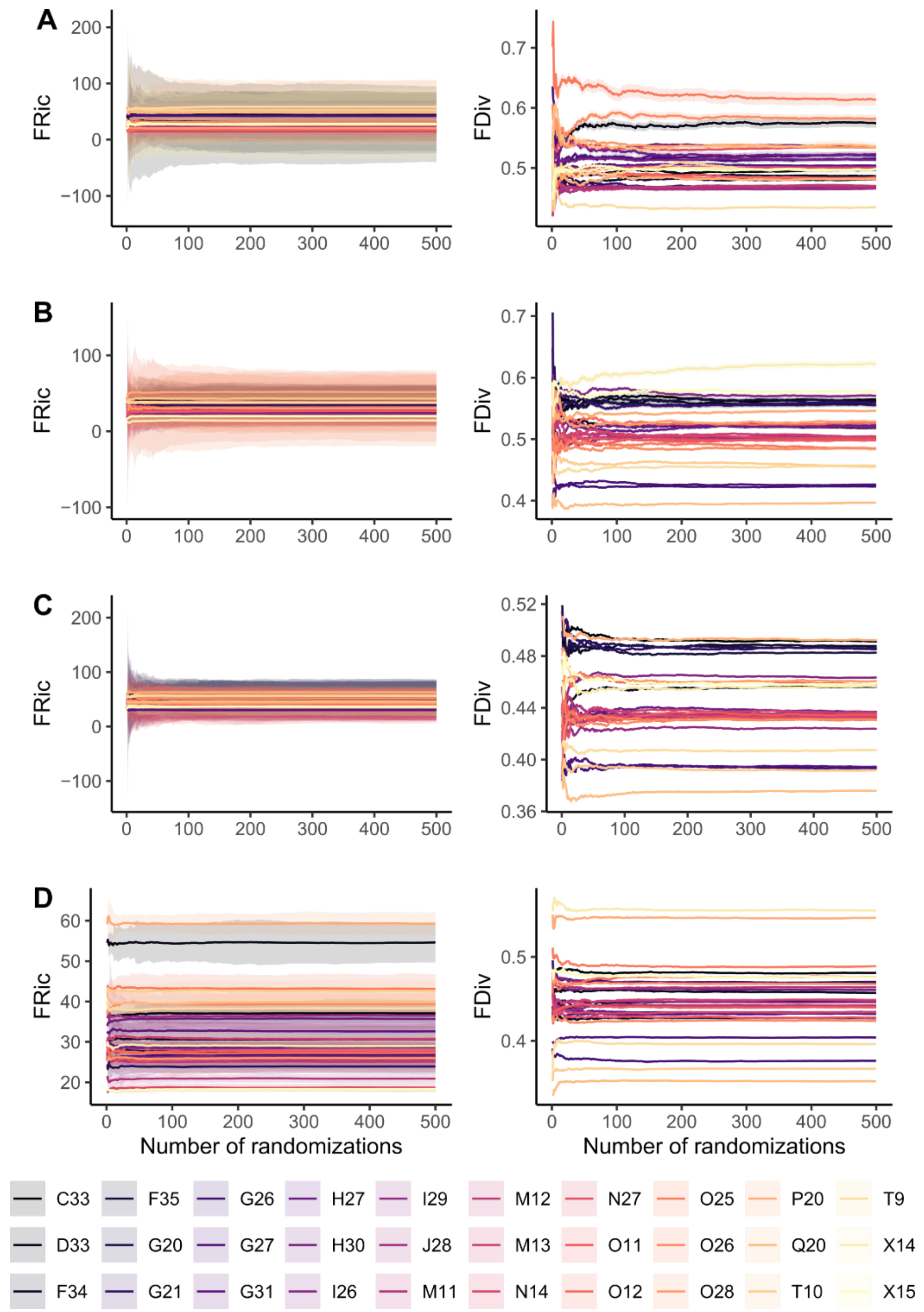


**Fig. S19. Evolution of mean and variance of simulated values of FRic and FDiv from different null models with increasing number of randomizations.** In order to assess the quality of using 500 randomizations, the changes in the mean (represented by lines) and the variance (represented as dashed areas) of functional diversity indices were studied in response to the number of randomizations for the null models of (**A**) random population, (**B**) random tree, (**C**) random leaf and (**D**) population-restricted random leaf null model. Colors correspond to the different plots included in our study.

**Table. S1. Summary of a principal component analysis for seven leaf functional traits, including loadings, standard deviation, proportion of the variance explained by each component and the adjusted eigenvalue obtained in a Horn's parallel analysis.**

| Trait | PC1 | PC2 | PC3 | PC4 | PC5 | PC6 | PC7 |
|---|---|---|---|---|---|---|---|
| SLA | 0.90 | 0.07 | -0.03 | -0.31 | -0.01 | 0.28 | -0.15 |
| LDMC | -0.94 | 0.19 | -0.07 | 0.16 | 0.08 | -0.06 | -0.22 |
| C | -0.73 | 0.29 | 0.19 | -0.44 | -0.38 | -0.03 | 0.02 |
| N | 0.46 | 0.75 | 0.16 | -0.21 | 0.28 | -0.29 | -0.01 |
| P | 0.38 | 0.64 | 0.46 | 0.40 | -0.25 | 0.09 | -0.01 |
| SDens | -0.64 | 0.63 | -0.18 | -0.03 | 0.24 | 0.32 | 0.09 |
| SSize | -0.32 | -0.42 | 0.81 | -0.09 | 0.24 | 0.09 | 0.00 |
| Standard deviation | 1.75 | 1.29 | 0.98 | 0.72 | 0.64 | 0.53 | 0.28 |
| Proportion of variance | 0.44 | 0.23 | 0.14 | 0.08 | 0.06 | 0.04 | 0.01 |
| Cumulative Proportion | 0.44 | 0.68 | 0.82 | 0.89 | 0.95 | 0.99 | 1.00 |
| Adjusted eigenvalue | 3.03 | 1.64 | 0.94 | 0.53 | 0.42 | 0.31 | 0.13 |

**Table. S2. Results for linear mixed-effects models studying the effects of tree species richness on multivariate functional indices used to estimate intraspecific variability, intraindividual variability and intraspecific overlap.** Estimates (standard errors) and significance assessed with likelihood ratio tests are shown. The slope of the terrain (slope) was included as a covariate in the models for intraspecific variability and intraspecific overlap, while the models for intraindividual variability included slope and diameter at breast height (DBH) as covariates.

| Level | Index | Tree species richness | Slope | DBH | $R^2$m | $R^2$c |
|---|---|---|---|---|---|---|
| Intraspecific | FRic | **-0.75 (0.36)*** | 0.07 (0.06) | - | 0.06 | 0.17 |
| Intraspecific | FDiv | -0.003 (0.004) | -0.0005 (0.001) | - | 0.04 | 0.11 |
| Intraindividual | FRic | -0.40 (0.68) | -0.03 (0.05) | **0.06 (0.02)*** | 0.03 | 0.29 |
| Intraindividual | FDiv | -0.001 (0.002) | 0.0001 (0.0003) | 0.0001 (0.0001) | 0.00 | 0.01 |
| - | Intraspecific overlap | **0.02 (0.01)*** | -0.002 (0.001) | - | 0.06 | 0.29 |

Note: $R^2$m, marginal $R^2$; $R^2$c, conditional $R^2$;

*$p < 0.05$.

**Table. S3. Species included in the study.** Species names and families from World Flora Online (https://www.worldfloraonline.org/; accessed 19 June 2024).

| Species | Family |
|---|---|
| *Castanea henryi* Rehder & E.H.Wilson | Fagaceae |
| *Castanopsis sclerophylla* (Lindl. & Paxton) Schottky | Fagaceae |
| *Choerospondias axillaris* (Roxb.) B.L.Burtt & A.W.Hill | Anacardiaceae |
| *Liquidambar formosana* Hance | Altingiaceae |
| *Nyssa sinensis* Oliv. | Nyssaceae |
| *Quercus serrata* Murray | Fagaceae |
| *Sapindus mukorossi* Gaertn. | Sapindaceae |
| *Triadica sebifera* (L.) Small | Euphorbiaceae |

**Table. S4. Results for linear mixed-effects models studying the effects of tree species richness and type of null model on standardized effect sizes of two functional indices.** Significance assessed with likelihood ratio tests are shown. The interaction between the predictors is indicated by ":".

| Response variable | Tree species richness | Type of null model | Tree species richness : Type of null model | $R^2m$ | $R^2c$ |
|---|---|---|---|---|---|
| $SES_{FRic}$ | *** | *** | *** | 0.26 | 0.61 |
| $SES_{FDiv}$ | *** | *** | *** | 0.40 | 0.68 |

Note: $R^2m$, marginal $R^2$; $R^2c$, conditional $R^2$;

***p < 0.01.

**Table. S5. Layers and hyperparameters used for building a convolutional neural network for every trait, and coefficient of determination (R2) and root mean squared error (RMSE) for the test and the train samples.**

| Layer | Hyperparameter | SLA | LDMC | C | N | P | SDens | SSize |
|---|---|---|---|---|---|---|---|---|
| Spectral region | - | 400-2500 | 400-2500 | 400-2500 | 1500-2400 | 1500-2400 | 400-2500 | 400-2500 |
| 1 dimension convolutional layer | Number of filters | 2 | 2 | 1 | 2 | 2 | 2 | 2 |
| 1 dimension convolutional layer | Kernel size | 50 | 2 | 35 | 77 | 77 | 2 | 1 |
| Batch normalization layer | - | Yes | Yes | Yes | Yes | No | Yes | No |
| Max-pooling layer | Pool size | 2 | 2 | 2 | 2 | 2 | 2 | 2 |
| Layer flatten | - | Yes | Yes | Yes | Yes | Yes | Yes | Yes |
| Layer dense | Number of nodes | 128 | 64 | 128 | 64 | 256 | 128 | 64 |
| Layer dense | Number of nodes | 32 | 16 | 32 | 16 | 64 | 32 | 16 |
| Layer dense | Number of nodes | 8 | 4 | 4 | 4 | 16 | 8 | 4 |
| - | Epochs | 500 | 2000 | 1000 | 1500 | 120 | 150 | 700 |
| - | Validation Split | 0.2 | 0.2 | 0.2 | 0.2 | 0.2 | 0.2 | 0.01 |
| $R^2$ test | - | 0.91 | 0.91 | 0.7 | 0.85 | 0.54 | 0.64 | 0.64 |
| RMSE test | - | 18.19 | 20.71 | 1.13 | 0.16 | 5.29 | 120.7 | 3.05 |
| $R^2$ train | - | 0.93 | 0.96 | 0.61 | 0.91 | 0.62 | 0.76 | 0.83 |
| RMSE train | - | 14.94 | 13.72 | 0.88 | 0.13 | 4.74 | 118.11 | 1.71 |

**Table. S6. Distribution of missing trait data in the leaf-level dataset across species and traits.**

| | SLA | LDMC | C | N | P | Sdens | Ssize | Mean |
|---|---|---|---|---|---|---|---|---|
| *Castanea henryi* | 3 (0.07%) | 3 (0.07%) | 92 (2.01%) | 22 (0.48%) | 30 (0.66%) | 79 (1.73%) | 41 (0.9%) | 38.57 (0.84%) |
| *Castanopsis sclerophylla* | 34 (0.74%) | 37 (0.81%) | 14 (0.31%) | 46 (1.01%) | 28 (0.61%) | 19 (0.42%) | 92 (2.01%) | 38.57 (0.84%) |
| *Choerospondias axillaris* | 0 (0%) | 0 (0%) | 23 (0.5%) | 11 (0.24%) | 28 (0.61%) | 1 (0.02%) | 5 (0.11%) | 9.71 (0.21%) |
| *Liquidambar formosana* | 4 (0.09%) | 3 (0.07%) | 5 (0.11%) | 32 (0.7%) | 33 (0.72%) | 14 (0.41%) | 22 (0.48%) | 16.14 (0.35%) |
| *Nyssa sinensis* | 15 (0.33%) | 21 (0.46%) | 4 (0.09%) | 4 (0.09%) | 12 (0.26%) | 2 (0.04) | 28 (0.61%) | 12.28 (0.27%) |
| *Quercus serrata* | 4 (0.09%) | 11 (0.24%) | 5 (0.11%) | 5 (0.11%) | 13 (0.28%) | 62 (1.36%) | 3 (0.07%) | 14.71 (0.32%) |
| *Sapindus mukorossi* | 1 (0.02%) | 2 (0.04%) | 0 (0%) | 31 (0.68%) | 66 (1.44%) | 1 (0.02%) | 7 (0.15%) | 15.43 (0.34%) |
| *Triadica sebifera* | 0 (0%) | 0 (0%) | 0 (0%) | 22 (0.48%) | 22 (0.48%) | 0 (0%) | 5 (0.11%) | 7 (0.15%) |
| Total | 61 (1.34%) | 77 (1.69%) | 143 (3.13%) | 173 (3.79%) | 221 (4.84%) | 178 (3.9%) | 203 (4.44%) | 150.86 (3.3%) |
